# Establishment and Resilience of Transplanted Gut Microbiota in Aged Mice

**DOI:** 10.1101/2021.03.18.435923

**Authors:** Ying Wang, Jinhui Tang, Qingqing Lv, Yuxiang Tan, Xiaoxiao Dong, Hongbin Liu, Nannan Zhao, Zhen He, Yan Kou, Yan Tan, Xin-an Liu, Liping Wang, Yang-Yu Liu, Lei Dai

**Author notes:** These authors contributed equally to this work (Ying Wang, Jinhui Tang).

## Abstract

Fecal microbiota transplantation (FMT), a procedure in which fecal material is transferred from a donor to a recipient, has been increasingly used as a treatment to restore healthy gut microbiota. There is a substantial difference in the composition of gut microbiota between young and aged hosts, but little is known about whether age matching between the FMT donor and recipient affects microbiota restoration and long-term maintenance. In the present investigation, we aimed to study the establishment and resilience of transplanted gut microbiota in aged recipients. We treated naturally aged mice (20 months old) with a broad-spectrum antibiotic cocktail and monitored the restoration of gut microbiota over 8 weeks. The diversity of gut microbiota in aged mice failed to reach the baseline level via spontaneous recovery; in contrast, FMT from either (age-)matched or unmatched donors facilitated the recovery of gut microbiota diversity. The microbiota transplanted from different donors successfully established in the aged recipients and had long-term effects on the gene expression profiles of the host colon. Finally, we evaluated the long-term maintenance of transplanted microbiota via intentional disruption of gut homeostasis. We found that lack of age matching between FMT donors and recipients may decrease the resilience of transplanted gut microbiota against colonic inflammation. The results from our study systematically examining the effects of FMT on the gut homeostasis of aged hosts suggest that the compatibility between donors and recipients should be taken into account when implementing FMT.

## Introduction

The gut microbiota is a complex and highly diverse ecosystem that maintains a symbiotic relationship with the host and regulates gastrointestinal (GI) homeostasis. Extensive studies have shown that the composition of gut microbiota is significantly different between the young and the aged in both humans and rodent models^1–3^. Alterations of the gut microbiome composition may be important regulators of host health. Many GI diseases, including *Clostridioides difficile* infection (CDI), inflammatory bowel disease, and irritable bowel syndrome are more prevalent in the elderly^4–6^. The probability of occurrence of complications and opportunistic infections were also higher in elderly patients^7,8^. The maintenance of healthy and resilient gut microbiota is critical for the life quality and healthspan of the elderly.

Fecal microbiota transplantation (FMT) is a procedure in which fecal material is transferred from a healthy donor to a patient with disrupted gut microbiota^9,10^. One recent study showed that autologous FMT improved the reconstitution of the post-antibiotic mucosal microbiome and gut transcriptome in humans and rodent models^11^. However, autologous FMT is not always a clinically available option, especially for aged hosts, and rational donor selection largely determines the patient response^12–15^. One study of ulcerative colitis patients reported that long-term post-FMT maintenance was significantly affected by the age difference between donors and patients^16^. Another study of FMT for CDI patients found a higher relapse rate among elderly patients compared with younger counterparts during the 6-month period after FMT^17^. These clinical observations underscore the issue of donor-recipient compatibility in FMT, as the age difference between recipients and donors may contribute to the long-term prognosis.

In this study, we used naturally aged mice (20 months old) as FMT recipients to study the establishment and resilience of transplanted gut microbiota. We treated aged mice with an antibiotic cocktail to disrupt the gut microbiota and monitored the recovery process either without FMT (spontaneous recovery) or with FMT (facilitated recovery). Compared with the spontaneous recovery (SR) group, we found that FMT facilitated the restoration of gut microbiota diversity and colon gene expression as the transplanted microbiota established in the new host. Furthermore, we tested how the transplanted microbiota responded to colon inflammation, which caused transient perturbations in gut homeostasis. We found that the gut microbiota transplanted from age-unmatched donors failed to withstand the perturbations, supporting the hypothesis that the resilience of transplanted microbiota is affected by the donor-recipient compatibility.

## Results

### Spontaneous recovery of gut microbiota following antibiotic treatment in aged mice

Previous studies have shown that antibiotic treatment can disrupt the gut microbiota in young hosts, leading to a substantial decrease in microbial load^18,19^ and microbiota diversity^11,20,21^. Afterwards, there is a long-term restorative process, during which the gut microbial ecosystem is temporarily imbalanced and significantly distinct from the original state^11,20–22^. However, little is known about the recoverability of gut microbiota in aged hosts.

To study the disruption of gut microbiota by antibiotics and the ensuing spontaneous recovery in aged hosts, we treated aged (20-month-old) female C57BL/6J mice with oral gavage of a broad-spectrum antibiotic cocktail (ampicillin, vancomycin, neomycin, and metronidazole) at a high dose (ABH group) or low dose (ABL group) or with water (Ctrl group). Fecal samples were collected before treatment and then regularly collected until 56 days after treatment (**Figure 1A**). We found that antibiotic treatment led to a substantial but transient decrease in bacterial load (the total copy number of the 16S rRNA gene) in feces (**Figure 1B**). In contrast, antibiotic treatment had a significant and long-term (8 weeks after treatment) impact on the number of observed species (**Figure 1C**) and the Shannon diversity index (**Figure 1D**) of gut microbiota.

**Figure 1.**
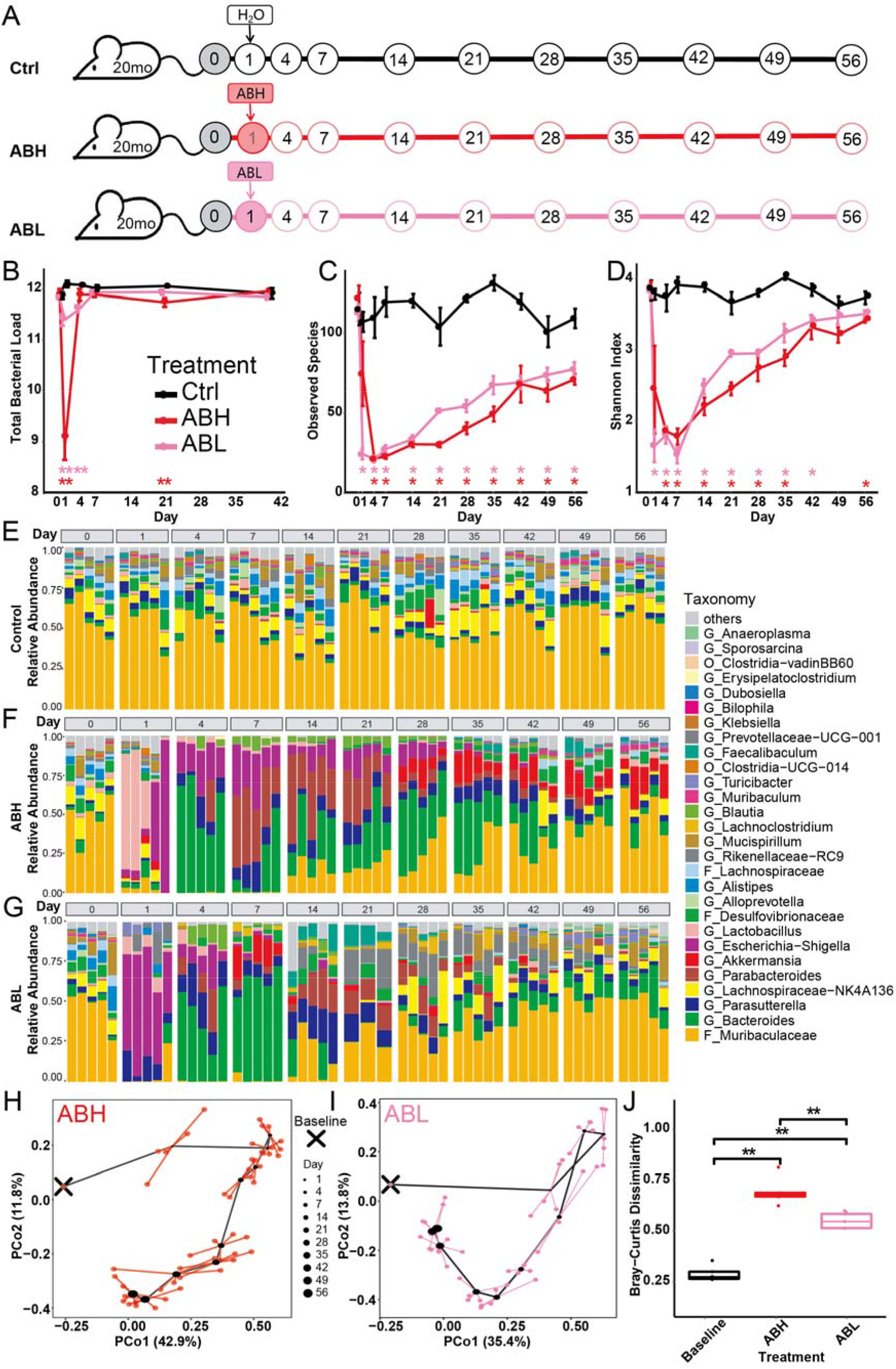
Spontaneous recovery of gut microbiota following antibiotic treatment in aged mice. (**A**) Aged mice were treated with the an antibiotic cocktail on Day 1. Ctrl, Control; ABH, antibiotics-high dose; ABL, antibiotics-low dose; 20mo: 20-month-old mice. Circles mark the time points (Day) of fecal sample collection. N=5 for each time point (except for the ABL group on Day 21, N=3). (**B**) Total microbial load (16S copy number log/gram) in fecal samples assayed by qPCR. (**C-D**) The number of observed species (**C**) and the Shannon diversity index (**D**) did not recover to the baseline level. Error bars= SEM. Colored asterisks indicate statistically significant differences between the experimental groups and the control group. (**E-G**) Changes in the relative abundance of microbial taxa during spontaneous recovery. O, order; F, family; G, genus. (**H-I**) Principal Coordinate Analysis (PCoA) based on Bray-Curtis dissimilarity, showing the trajectory of microbiota composition for the ABH (**H**) and ABL (**I**) groups. Baseline (black cross) indicates the microbiota composition averaged over aged mice in the control group from Day 1 to Day 56. For each time point, the central black dot indicates the average of five mice and the colored lines connect replicate samples. (**J**) Bray-Curtis dissimilarity between the gut microbiota composition on Day 56 and the baseline composition. The FDR-adjusted P values are shown between various treatment groups (two-sided Wilcoxon rank sum test). \**P*< 0.05, \*\**P*< 0.01, \*\*\**P*< 0.001.

The alpha diversity of gut microbiota in aged mice did not recover to the baseline level, suggesting the elimination of community members during the antibiotic treatment^21^. These observations are consistent with previous studies in young mice^11^ and in an independent experiment that we carried out in 2-month-old mice (**Figure S1**).

We found that the composition of gut microbiota in aged mice was disrupted by antibiotic treatment and gradually recovered to a diverse community over 8 weeks (**Figure 1E–1G**). The microbiota composition of the control group (i.e., no antibiotic treatment) remained stable throughout the entire period (**Figure 1E**), while the gut microbiota in the antibiotic treatment groups (ABH and ABL) changed dramatically during the first 2 weeks. We observed a clear pattern of ecological succession in the absolute abundance profiles (**Figure S2**): some bacterial taxa transiently flourished and dominated the microbial community and then were replaced by other taxa. Consistent with previous studies on the post-antibiotic recovery of gut microbiota in young mice and humans^23–25^, we observed some “recovery-associated bacterial species”, such as *Bacteroides* and *Alistipes*, which increased in abundance early during recovery. After Day 14, the relative abundance of most bacterial taxa stabilized, but some taxa (e.g., *Muribaculaceae*) took longer to return to their baseline level. Similar to the findings in young mice (^11^ and **Figure S1**), we found that the gut microbiota in aged mice had not returned to the baseline composition by 8 weeks after the antibiotic treatment (**Figure 1H and 1I**). On Day 56, the gut microbiota compositions of the treatment groups (ABH, ABL) were significantly different from the baseline composition (**Figure 1J**).

### Establishment of transplanted gut microbiota in aged mice

Since heterochronic FMT is usually the only option for elderly patients in clinical situations, it is essential to investigate whether FMT donors with significantly different microbiota are equally successful in facilitating microbiota restoration in recipients. To verify whether donor-recipient compatibility affects the outcome, two parallel groups of aged mice were given a high-dose antibiotic cocktail treatment and then received FMT from age-matched 20-month-old donor mice (FMT-M), or aged-unmatched 2-month-old donor mice (FMT-UM). FMT was performed by oral gavage for three consecutive days after antibiotics treatment (**Figure 2A and Methods**). We found that the two groups of donor mice (2 months old vs. 20 months old) had very distinct microbiota (**Figure S3A and S3B**). Differential abundance analysis using LEfSe (Linear discriminant analysis Effect Size) revealed that *Prevotellaceae*and *Bilophila* were enriched in the baseline microbiota of young mice, whereas *Bacteroides*and *Desulfovibrio* were enriched in aged mice (**Figure S3C**).

**Figure 2.**
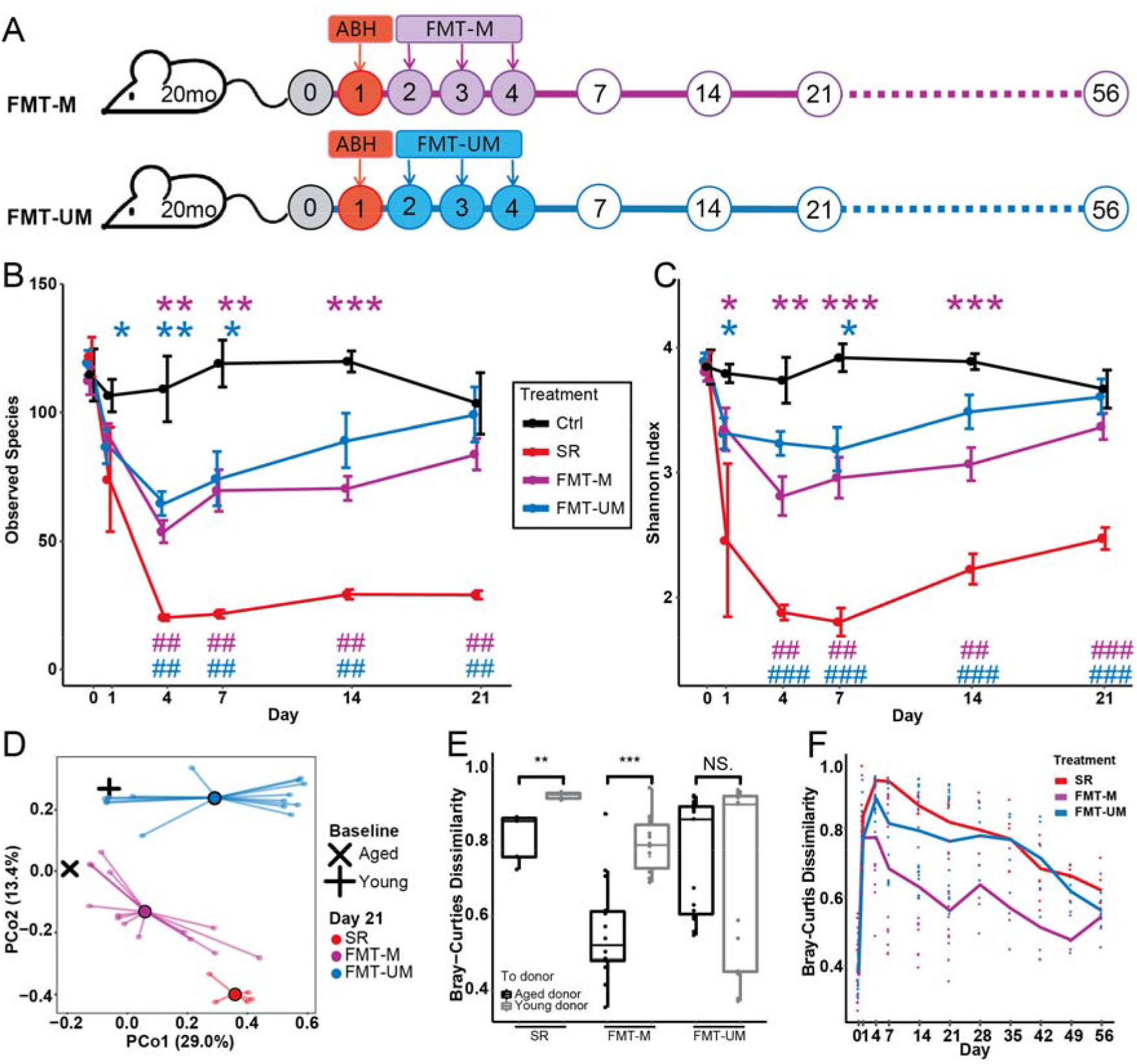
Establishment of transplanted gut microbiota in aged mice. (**A**) Following high-dose antibiotics treatment, FMT was performed for aged mice between Day 2 and Day 4 using age-matched and age-unmatched donors. Ctrl, Control. SR, spontaneous recovery. FMT-M, FMT using age-matched donors. FMT-UM, FMT using age-unmatched donors. (**B-C**) FMT facilitates the restoration of gut microbiota diversity in aged mice. In contrast to the SR group, the number of observed species (**B**) and Shannon diversity index (**C**) of the gut microbiota of the FMT groups returned to the baseline level in 3 weeks. Error bars= SEM.* indicates significant difference at *P*< 0.05 compared with the Ctrl group on the same day; # indicates significant difference at P < 0.05 compared with the SR group on the same day. **or ##, *P*<0.01, *** or ###, *P*<0.001. (**D**) The post-FMT gut microbiota on Day 21 moved toward the baseline composition of the donors (FMT-M: aged donor, FMT-UM, young donor). The baseline microbiota composition is averaged over 30 aged mice (20 months old) or young mice (2 months old) on Day 0. The central black dot indicates the average of 5 mice (SR group) or 15 mice (FMT-M and FMT-UM groups) and the colored lines connect replicate samples. (**E**) Bray-Curtis dissimilarity between the post-FMT gut microbiota on Day 21 and the baseline microbiota (i.e., aged donors or young donors). (**F**) Bray-Curtis dissimilarity between the gut microbiota of treatment groups and the baseline microbiota of aged mice throughout the long-term recovery. For the FMT-M and FMT-UM groups, N=15 until Day 21 and N=5 thereafter. N=5 for the Ctrl and SR groups. The FDR-adjusted P values are shown between various treatment groups (two-sided Wilcoxon rank sum test).

The change in the number of observed species (**Figure 2B**) and Shannon index (**Figure 2C**) showed that FMT substantially accelerated the recovery of gut microbiota diversity in comparison with spontaneous recovery. The gut microbiota diversity of aged mice that received FMT was fully restored by Day 21 after antibiotic treatment (**Figure 2B and 2C**). This is consistent with a previous study, which showed that autologous FMT improved the post-antibiotic reconstitution of the gut microbiome in young hosts (mice and human volunteers)^11^.

On Day 21, the microbiota composition of mice that received FMT was more similar to the baseline of young or aged mice (average microbiota composition of donors on Day 0), in comparison to mice in the SR group (**Figure 2D**). In addition, the gut microbiota of mice in the FMT-M group was closer to the baseline microbiota of aged donors, while there was larger variation in the gut microbiota of mice in the FMT-UM group, with one sub-group having a microbiota close to the baseline of young donors (**Figure 2E**). Kinetics variation of microbiota composition recovery after antibiotic treatment has been observed in previous studies in conventional and humanized young mice^21^. However, our results revealed a novel finding that aged mice in the FMT-M group had more consistent compositional kinetics than those in the FMT-UM group. The individual differences in gut microbial composition on Day 21 between mice in both the FMT-M and FMT-UM groups were not fully attributable to cage effects (**Figure S4**).

We demonstrated that age-unmatched donors were as effective as age-matched donors in restoring the diversity of the microbiota, with both groups showing a significant increase in diversity when compared with the SR group. Moreover, the composition of newly established microbiota was donor dependent. We followed five recipient mice in each FMT group until Day 56 (**Figure S5**). For each mouse, we calculated the Bray-Curtis dissimilarity between its gut microbiota at different time points and the original baseline gut microbiota (i.e., Day 0). We found that the gut microbiota of mice in the FMT-M group was closer to the baseline of aged mice than the other two groups throughout the entire experiment (**Figure 2F and Figure S5A**).

### Long-term effects of FMT on the gut metagenome and colon gene expression

Previous research has shown that that functional restoration of bacteriomes and viromes revealed by metagenomic sequencing could be an indicator of successful FMT^26^. To evaluate the long-term effects of FMT in aged mice with different donors, we performed shotgun metagenomic sequencing of Day-56 fecal samples from each treatment group (SR, FMT-M, and FMT-UM) together with the baseline of 20-month-old mice (matched donors, MD) and 2-month-old mice (unmatched donors, UMD). We found that 35 out of 219 functional pathways were differentially expressed between the SR group and MD (**Figure 3A**). We then examined the levels of these pathways in the FMT-M and FMT-UM groups after 56 days of recovery. We found that most of the 35 pathways that did not spontaneously recover to baseline levels were restored in the FMT-M group. In contrast, FMT-UM was not as effective in facilitating recovery of microbiota function. When all 219 functional pathways were taken into consideration, endpoint samples of both the FMT-M and FMT-UM groups were closer to the baseline samples compared with samples from the SR group on Day 56 (**Figure S6**).

**Figure 3.**
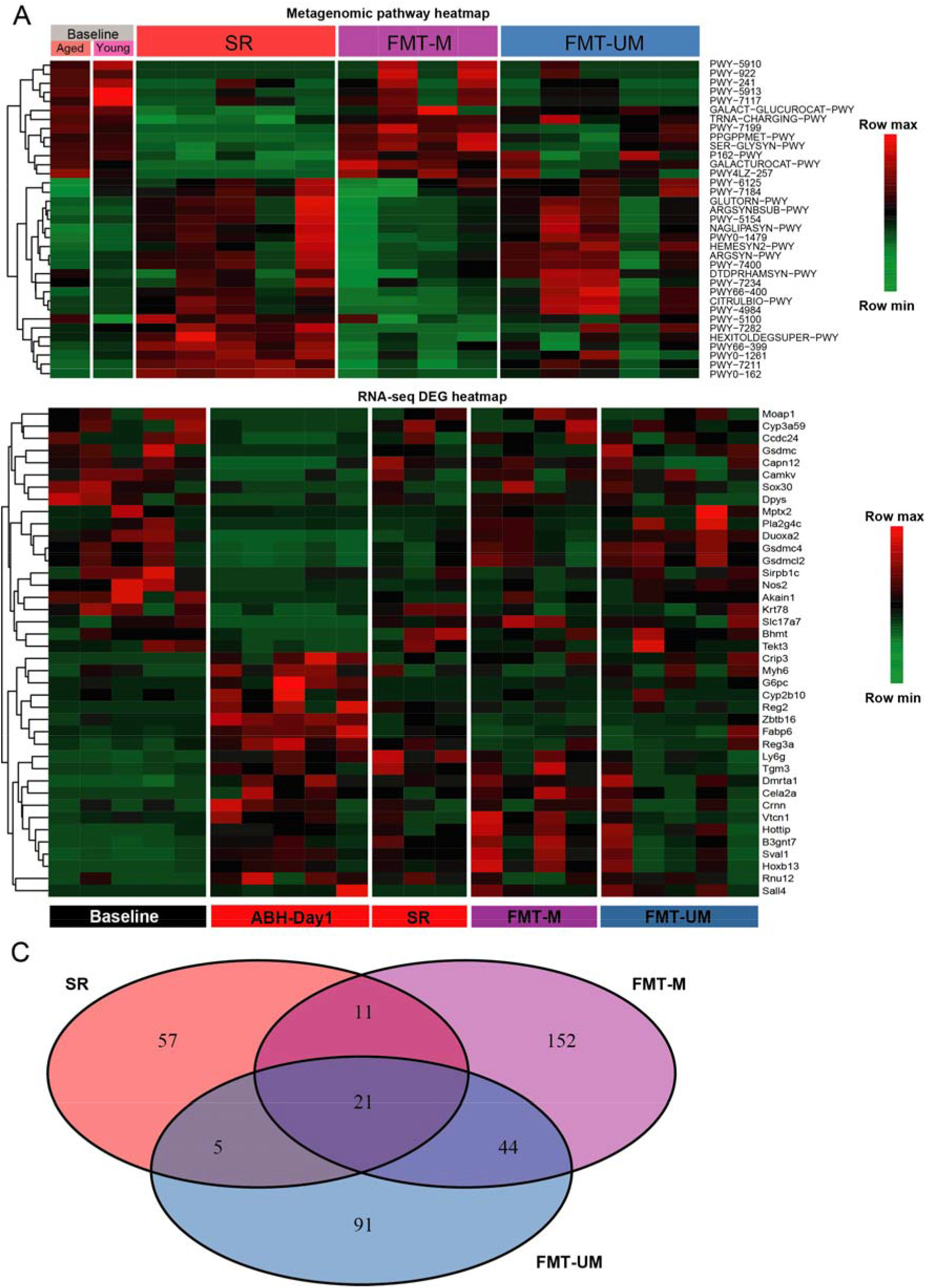
Long-term effects of FMT on the gut metagenome and colon gene expression in aged mice. (**A**) Metagenomic gene pathways that were differentially abundant (FDR-corrected *P*<0.01) between the SR group on Day 56 and the baseline of aged mice. SR, spontaneous recovery. FMT-M, FMT using age-matched donors. FMT-UM, FMT using age-unmatched donors. SR/FMT-M/FMT-UM samples were collected on Day 56. (**B**) Heatmap of differentially expressed genes (DEGs) (*P*< 0.05) between ABH-Day1 and Baseline, based on RNA-seq of colon samples. Baseline, aged mice on Day 0. ABH-Day1, samples collected on Day 1 after high-dose antibiotic treatment. SR/FMT-M/FMT-UM samples were collected on Day 56. (**C**) Venn diagram showing the DEGs (*P*< 0.05) between the baseline and treatment groups. Most of the DEGs were present in only one group.

The shift in the transcriptome of the host following antibiotics treatment can be dramatic^11,23,27^, and the response is determined by the combined effects of microbiota depletion and the remaining antibiotic-resistant microbes as well as the direct effects of antibiotics on host tissues^27^. It was shown that autologous FMT in young mice helped restore the gene expression profile across GI tracts to some extent^11^. However, the effects of FMT donor on the transcriptional profiles of aged hosts remain unknown. We performed RNA sequencing (RNA-seq) of upper colon samples of aged mice before treatment (Baseline), Day 1 after antibiotics (ABH-Day1), and at the end point (Day 56) for the three treatment groups (SR, FMT-M, and FMT-UM). We first examined the restoration of differentially expressed genes (DEGs) between ABH-Day1 and the Baseline on Day 56 in the SR, FMT-M, and FMT-UM groups (**Figure 3B**). Most of the genes that were affected by antibiotics treatment were not reverted to the Baseline configuration in either the SR group or the FMT groups (**Figure 3B**). We then examined the DEGs between each group on Day 56 and the Baseline and the overlap among them (**Figure 3C and Figure S7**). Interestingly, a large fraction of DEGs were not shared between the different treatment groups (SR, FMT-M, and FMT-UM), which demonstrated that the transcriptional landscapes on Day 56 were highly dependent on FMT and the choice of donor.

### The resilience of transplanted gut microbiota against colon inflammation

Whether the transplanted microbiota is resilient against perturbations to gut homeostasis reflects its long-term stability. We utilized a self-limiting colon inflammation model induced by administration of 3% dextran sulfate sodium (DSS) in the drinking water of the mice for 7 days, which is widely used to induce acute colitis in rodents^28,29^. Previous studies had shown that DSS-induced-colitis caused changes in the gut microbiota composition, which were spontaneously restored to the original state after the end of DSS treatment^30–32^.

In the sections above we showed that the microbiota diversity returned to baseline levels on Day 21 in the FMT-M and FMT-UM groups (**Figure 2B and 2C**). Also, microbial compositions largely stabilized after Day 21 (**Figure S5**). Hence, we chose Day 26 as the starting point for DSS treatment. From Day 26 to Day 31 after antibiotic treatment, aged recipients of FMT-M or FMT-UM were given 3% DSS treatment for seven consecutive days and then switched back to normal drinking water for 42 days (**Figure 4A**). Aged recipient mice of FMT-M or FMT-UM both showed typical symptoms of acute colitis as shown by weight loss (**Figure S8A**) and diarrhea and blood in feces (**Figure S8B**). These disease symptoms slowly resolved after the mice were switched back to normal drinking water, which is the same as what was reported in young and non-transplanted mice^33–35^.

**Figure 4.**
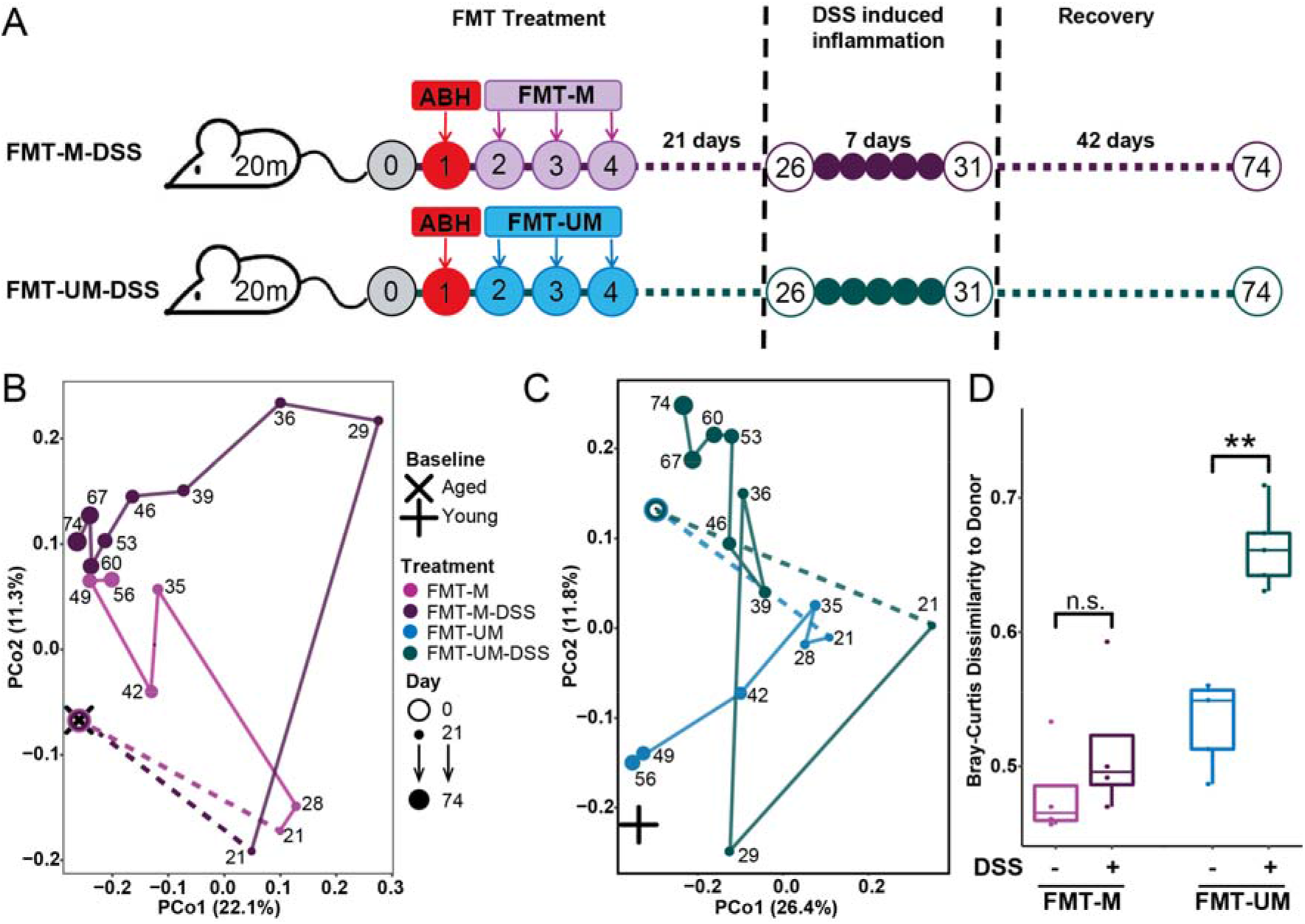
The resilience of transplanted gut microbiota against colon inflammation in aged mice. (**A**) Three weeks after FMT treatment (Day 26), aged mice were treated with DSS for 7 days to induce acute colon inflammation. Fecal samples were collected every week until Day 74. (**B-C**) PCoA analysis based on Bray-Curtis dissimilarity of microbiota composition at different time points. The comparison between the FMT-M group (FMT from aged-matched donor, without DSS perturbation) and FMT-M-DSS group (FMT from aged-matched donor, with DSS perturbation) are shown in (**B**). Similarly, the comparison between the FMT-UM and FMT-UM-DSS groups is shown in (**C**). For each group, the dot is the average of five mice (N=5) at each time point. (**D**) Bray-Curtis dissimilarity between the final microbiota composition of treatment groups and the baseline microbiota of the respective FMT donors (aged baseline: FMT-M and FMT-M-DSS; young baseline: FMT-UM and FMT-UM-DSS). Central line, median; box limits, first and third quartile; whiskers, 1.5 X interquartile range. The FDR-adjusted P values are shown between linked treatment groups (two-sided Wilcoxon rank sum test). * indicates significant difference at *P*< 0.05, **,*P*<0.01, ***, *P*<0.001.

PCoA based on Bray-Curtis dissimilarity was used to compare the changes in microbiota composition in FMT recipients with or without DSS treatment (**Figure 4B and 4C**). For mice in the FMT-M-DSS group, the microbiota restoration process was only briefly disrupted by the induced inflammation; the microbiota composition after Day 53 was similar to that of FMT-M group (without DSS treatment) (**Figure 4B**). At the end point of the experiments, the microbiota composition of FMT-M mice and that of FMT-M-DSS mice showed no significant difference in the distance to that of their aged donors (**Figure 4D**).

In contrast, for mice in the FMT-UM-DSS group, the microbiota composition after DSS treatment gradually shifted toward the original baseline of aged mice (**Figure 4C**). At the end point, compared with the FMT-UM mice, the microbiota composition of FMT-UM-DSS mice was significantly further from the baseline of its young donors (**Figure 4D**). These observations suggested that under the conditions of colon inflammation, the transplanted gut microbiota from age-unmatched donors may not be as stable as that from age-matched donors.

## Discussion

Heterologous FMT is the most widely used gut microbiome intervention in clinical settings. Most research so far has focused on the efficacy and safety of FMT^36–40^. Several studies reported that heterochronic FMT could result in the transfer of some physiological properties from the donor to the recipient, including lifespan and healthspan^41,42^, the germinal center reaction^43^, and neurogenesis and behavior^44,45^. However, less attention has been paid to the “pharmacokinetics” of FMT. Although FMT has become the most effective treatment for recurrent CDI, it was reported that about 20% of patients still experience recurrent CDI after the initial FMT therapy^46,47^. Previous studies investigating FMT treatment for IBD patients also suggested that a second or multiple FMT procedures are needed for better clinical outcomes and that the timing of the subsequent FMT is critical in determining the treatment effects^48,49^. These results suggest that monitoring the long-term effects of FMT and identifying the determinants of relapse after FMT are critical in making rational clinical decisions.

In this study, we focused on the establishment and resilience of transplanted gut microbiota from different donors by following the post-FMT dynamics in aged mice for 8 weeks. One intriguing observation was that recipients of transplanted microbiota from age-unmatched donors could not maintain the donor microbiota composition after perturbations in gut homeostasis, suggesting that the choice of donor may affect the resilience of gut microbiota after FMT. While studies on the time of maintaining donor-derived microbiota composition are emerging^48–50^, the resilience of the transplanted microbiota has less been studied. A resilient gut microbiota protects its host from both dysbiosis-related GI disorders and many non-GI diseases^51,52^. If the transplanted microbiota from unmatched donors is not resilient against perturbations, we should monitor the post-FMT dynamics of the gut microbiome and perform multiple FMT procedures if relapse occurs.

The mechanism that underlies the relationship between donor-recipient age-matching and the resilience of transplanted microbiota is still unknown. However, shaping of the microbiota by the aged host may play an important role. The microbiota and the host establish homeostasis through lifelong interactions, resulting in such a highly compatible state that the composition of gut microbiota may serve as a biomarker of aging^53,54^. When this highly compatible microbiota is suddenly replaced by another with a different composition through age-unmatched FMT, it may be in a metastable state and thus the balance can be more easily tipped by further disruptions^51^. Host immune status^55–57^ and metabolism^58^ may affect the resilience of transplanted microbiota. For example, innate immune surveillance systems co-evolve with the commensal microbiota-derived Toll-like receptors and Nucleotide oligomerization domain ligands, resulting in highly selective recognition and reactivity pattern in response to invasion by foreign species^56,57^. Transplanted microbiota from age-unmatched donors may trigger different immune responses, which in turn affect the microbiota resilience. Further animal and clinical studies are definitely needed to validate and understand the influences of hosts on the resilience of microbiota.

In our study and previous clinical reports^16^, the difference in the resilience of transplanted microbiota was attributed to the age difference between FMT donors and recipients. It is possible that age-matching is not the only factor determining the long-term stability of transplanted microbiota. Age-matched populations have very diverse gut microbiota due to differences in geographical distribution, diet, lifestyle, and many other factors. On the other hand, selection of donors from among family members, though age-unmatched, might result in a more similar microbiota composition than that obtained using aged-matched donors. The effect of FMT using age-matched but composition-unmatched donors (or vice versa) remains to be studied.

In summary, our present investigation sheds light on the long-term dynamics of the gut microbiome in aged hosts and suggests that donor-recipient compatibility should be taken into consideration during the implementation of FMT.

## Methods

### Animal experiments

16-month-old C57BL/6J mice were purchased from The Model Organisms Center, Inc. (Shanghai, CN) and naturally aged to 20-month-old in specific pathogen-free vivarium at the Shenzhen Institutes of Advanced Technology (SIAT). 2-month-old C57BL/6J mice were purchased from The Vital River Laboratories (Beijing, CN). All mice used in the present study were females. Mice were housed according to treatment groups. All animal experiments were approved by the Institutional Animal Care and Use Committee at SIAT. Fecal pellets were collected and snap-frozen in liquid nitrogen and transferred to storage at −80C. Upon the termination of experiments, mice were sacrificed by isoflurane inhalation. The upper colon (between 1/4 to 1/2 of the total length from beneath the cecum to the rectal) was dissected and stored in liquid nitrogen to be used for RNA sequencing.

### Antibiotic treatment

High-dose antibiotics cocktail (ABH) was prepared in sterile ddH_2_O at the following concentration: ampicillin (1 mg/mL), vancomycin (5 mg/mL), neomycin trisulfate (10 mg/mL), metronidazole (10 mg/mL). All antibiotics purchased form Sangon biotec, Shanghai. Low-dose antibiotics cocktail (ABL) was prepared by 10-fold dilution of ABH solution in ddH_2_O. Antibiotics cocktails were freshly prepared on the day of treatment and was administered by one-time oral gavage (250μL per mice) on Day 0 after the collection of fecal samples.

### Fecal microbiota transplantation

Fecal pellets were collected from age-matched donor mice (20-month-old) or aged-unmatched donor mice (2-month-old). Fecal pellets were pooled, mixed, and homogenized in PBS at concentration of 1g feces/10mL PBS^44,58^. Mixture was centrifuged at 500 rpm for 5 minutes at 4°C and the supernatant were collected and used for FMT. Each aged recipient mouse received 150ul of the supernatant by oral gavage once a day continuously for 3 days. For FMT with matched donors (FMT-M), the same group of mice were used as donor and recipients by collecting feces samples before antibiotics treatment to prepare FMT supernatant.

### DSS-induced colon inflammation

3% w/v dextran sulfate sodium (DSS) solution was prepared in autoclaved ddH_2_O and used to replace normal drinking water for 7 days. Mice had free access to water and the amount of water consumption was monitored to confirm that there was no difference among treatment groups. DSS solution was freshly made and changed for the mouse cages every 2-3 days.

Weight of each mice were measured at fixed time (9 am) everyday. Disease activity index (DAI) were calculated as a combined score of weight loss (0 or positive=0, 0.1-5%=1, 5-10%=2, 10-15%=3, >15%=4), stool consistency (normal=0, partially loose=1, loose stool=2, mild diarrhea=3, diarrhea=4), and rectal bleeding (normal=0, dark stool with no fresh blood=1, fresh blood seen in <50% pallets =2, blood regularly seen= 3, active bleeding= 4).

### Bacterial DNA extraction and qPCR

DNA was isolated from fecal samples using the QIAamp PowerFecal DNA Kit (Qiagen) according to manufacturer’s instructions. Using DNA extracted from fecal samples of 2-month-old mice, the 16S rRNA gene was amplified by primers 27F 5’-AGAGTTTGATCCTGGCTCAG-3’ and 1492R 5’-TACGGTTACCTTGTTACGACTT-3’ with PrimeSTAR^®^ Max DNA Polymerase and purified by gel extraction. The purified 16S rRNA gene is used as a standard in qPCR. The copy number of 16S rRNA gene in fecal samples was determined by qPCR using the TB Green Premix Ex Taq II (Tli RNaseH Plus) with primers 764F 5’-CAAACAGGATTAGATACCC-3’ and 907R 5’-CCGTCAATTCCTTTRAGTTT-3’^59^.

### 16S rRNA amplicon sequencing and metagenomic sequencing

V3-V4 region of 16S rRNA gene was amplified using primers 338F 5’-ACTCCTACGGGAGGCAGCA-3’ and 806R 5’-GGACTACCAGGGTATCTAAT-3’^59^ with 12bp barcode. PCR products were mixed in equidensity ratios. Then, mixture PCR products were purified with EZNA Gel Extraction Kit (Omega, USA). Sequencing libraries were generated using NEBNext Ultra DNA Library Prep Kit for Illumina (New England Biolabs, USA) and sequenced on an Illumina Hiseq2500 platform (250 bp paired-end reads), conducted by Guangdong Magigene Biotechnology Co.,Ltd. (Guangzhou, China). The average sequencing depth of each sample was around 70,000 reads. For metagenomic sequencing, DNA extracted from fecal samples (100 ng) was sheared with a Covaris E220X sonicator. Sequencing libraries were generated using NEB Next Ultra DNA Library Prep Kit for Illumina (New England Biolabs, USA) and sequenced on an Illumina Hiseq X-ten platform (150 bp paired-end reads), conducted by Guangdong Magigene Biotechnology Co.,Ltd. (Guangzhou, China).

### RNA-seq of colon samples

Approximately 50 mg were used for RNA isolation. Frozen colon tissues were homogenized, and RNA was isolated using TRIzol Reagent (Invitrogen). A total amount of 1 μg RNA per sample was used as input material for the library preparations. Sequencing libraries were generated using NEBNext UltraTM RNA Library Prep Kit for Illumina^®^ (NEB, USA). The clustering of the index-coded samples was performed on a cBot Cluster Generation System using TruSeq PE Cluster Kit v3-cBot-HS (Illumina). After cluster generation, the library preparations were sequenced on an Illumina Novaseq platform and 150 bp paired-end reads were generated. RNA extraction, library generation and sequencing were conducted by Novogene Co., Ltd. (Tianjin, China).

### Analysis of 16S rRNA amplicon sequencing data

16S sequencing data were analyzed by QIIME2 (version 2019.7)^60^. Primers of the raw sequence data were cut with Cutadapt (via q2-cutadapt)^61^, followed by denoising, merging and removing chimera with DADA2 (via q2-dada2)^62^. All amplicon sequence variants (ASVs) from DADA2 were used to construct a phylogenic tree with fasttree2 (via q2-phylogeny)^63^. The ASVs were assigned to taxonomy with naïve_bayes classifier (via q2-feature-classifier)^64^ against the SILVA database (SILVA_138_SSURef_Nr99). The ASV table was normalized by the sample with the fewest sequence reads, and rare ASVs (<0.1% in relative abundance) were filtered out. Alpha diversity was calculated by two measures, the Observed species based on the number of ASV and the Shannon diversity index per sample. Bray-Curtis dissimilarity and weighted unifrac distance was calculated by the Vegan package in R (https://CRAN.R-project.org/package=vegan). In stacking column plots, relative abundance was calculated to the lowest taxonomy level. If some reads from the same family were annotated at genus level, calculating these reads into family level. LEfSe^65^ was used to identify microbial taxa with differential abundant across groups.

### Analysis of metagenomics sequencing data

Raw reads were first trimmed using Trimmomatic v.0.39 (parameter setting: SLIDINGWINDOW:4:20, MINLEN:50)^66^, and then reads mapped to the host genome (mouse_C57BL_6NJ) or PhiX were filtered out using Bowtie2 (parameter setting: -very-sensitive-dovetail). Kraken2 v2.0.9-beta (with default parameters) was used to obtain the taxonomic classification of reads^67^. HUMAnN2 (with default parameters) was used to profile functional gene pathways with MetaCyc annotations^68^.

### Analysis of RNA-seq data

Raw reads were trimmed using Trimmomatic v.0.39 (parameter setting: SLIDINGWINDOW:4:20, MINLEN:50) and mapped to GRCm38 genome using STAR v2.7.4a (default parameters)^69^. Gene-level read counts were generated with featureCounts and Genecode annotation (GRCm38.p6.genome, gencode.vM25.annotationgtf). Genes with a minimum of 5 reads in at least one sample were included for downstream analysis. Normalization of the gene counts and differential expression analysis was performed by DESeq2^70^ to find differentially expressed genes. Statistical significance was assessed using a negative binomial Wald test, and then corrected for multiple hypothesis testing with the Benjamini-Hochberg method.

### Statistical analysis

Statistical analysis was performed in R (version 3.6.1). Statistically significant findings were marked according to the following cutoffs: *, *P*< 0.05; **, *P*< 0.01; ***, *P*< 0.001. Data were plotted with R ggplot2 package^71^. To account for multiple comparisons at each day, we considered a false discovery rate (FDR)-adjusted *P* value. Sample size for each experiment and the statistical tests used are specified in figure legends.

### Sequencing data availability

All sequencing data are deposited to NCBI Sequencing Read Archive (BioProject accession number PRJNA687886).

### Code availability

All scripts will be available on Github.

## Acknowledgments

We thank members of the LD lab for insightful discussions. Y Wang is supported by National Natural Science Foundation of China (31900839) and China Postdoctoral Science Foundation (2020M672880). L. Dai is supported by National Key R&D Program of China (2019YFA09006700) and National Natural Science Foundation of China (31971513). Y-Y Liu is supported by grants R01AI141529, R01HD093761, R01AG067744, UH3OD023268, U19AI095219, and U01HL089856 from National Institutes of Health, USA.

## Author Contributions

Y. Wang and L. Dai designed the research. Y. Wang, J. Tang, Q. Lv, X. Dong and H. Liu performed the experiments. J. Tang, Y. Wang, Y. Tan, X. Dong, N. Zhao, Y-Y Liu and L. Dai analyzed the data. J. Tang and Y. Wang made the figures. Y. Wang, Y-Y Liu and L. Dai wrote the manuscript. All authors critically revised and approved the final version.

## References

1 Claesson, M. J. et al. Composition, variability, and temporal stability of the intestinal microbiota of the elderly. Proc Natl Acad Sci USA 108 Suppl 1, 4586–4591, doi:10.1073/pnas.1000097107 (2011).

2 Jeffery, I. B., Lynch, D. B. & O’Toole, P. W. Composition and temporal stability of the gut microbiota in older persons. Isme j 10, 170–182, doi:10.1038/ismej.2015.88 (2016).

3 Thevaranjan, N. et al. Age-Associated microbial dysbiosis promotes intestinal permeability, systemic inflammation, and macrophage dysfunction. Cell Host Microbe 21, 455–466.e454, doi:10.1016/j.chom.2017.03.002 (2017).

4 Asempa, T. E. & Nicolau, D. P. Clostridium difficile infection in the elderly: an update on management. Clin Interv Aging 12, 1799–1809, doi:10.2147/cia.S149089 (2017).

5 Durazzo, M., Campion, D., Fagoonee, S. & Pellicano, R. Gastrointestinal tract disorders in the elderly. Minerva Med 108, 575–591, doi:10.23736/s0026-4806.17.05417-9 (2017).

6 Taleban, S., Colombel, J. F., Mohler, M. J. & Fain, M. J. Inflammatory bowel disease and the elderly: a review. J Crohns Colitis 9, 507–515, doi:10.1093/ecco-jcc/jjv059 (2015).

7 Lin, E., Lin, K. & Katz, S. Serious and opportunistic infections in elderly patients with inflammatory bowel disease. Gastroenterol Hepatol (N Y) 15, 593–605 (2019).

8 Bollegala, N., Jackson, T. D. & Nguyen, G. C. Increased postoperative mortality and complications among elderly patients with inflammatory bowel diseases: an analysis of the national surgical quality improvement program cohort. Clin Gastroenterol Hepatol 14, 1274–1281, doi:10.1016/j.cgh.2015.11.012 (2016).

9 Smillie, C. S. et al. Strain tracking reveals the determinants of bacterial engraftment in the human gut following fecal microbiota transplantation. Cell Host Microbe 23, 229–240.e225, doi:10.1016/j.chom.2018.01.003 (2018).

10 Zhang, F. et al. Microbiota transplantation: concept, methodology and strategy for its modernization. Protein Cell 9, 462–473, doi:10.1007/s13238-018-0541-8 (2018).

11 Suez, J. et al. Post-antibiotic gut mucosal microbiome reconstitution is impaired by probiotics and improved by autologous FMT. Cell 174, 1406–1423.e1416, doi:10.1016/j.cell.2018.08.047 (2018).

12 Vermeire, S. et al. Donor species richness determines faecal microbiota transplantation success in inflammatory bowel disease. J Crohns Colitis 10, 387–394, doi:10.1093/ecco-jcc/jjv203 (2016).

13 Wilson, B. C., Vatanen, T., Cutfield, W. S. & O’Sullivan, J. M. The super-donor phenomenon in fecal microbiota transplantation. Front Cell Infect Microbiol 9, 2, doi:10.3389/fcimb.2019.00002 (2019).

14 Kassam, Z. et al. Donor screening for fecal microbiota transplantation. N Engl J Med 381, 2070–2072, doi:10.1056/NEJMc1913670 (2019).

15 Duvallet, C. et al. Framework for rational donor selection in fecal microbiota transplant clinical trials. PLoS One 14, e0222881, doi:10.1371/journal.pone.0222881 (2019).

16 Okahara, K. et al. Matching between donors and ulcerative colitis patients is important for long-term maintenance after fecal microbiota transplantation. J Clin Med 9, doi:10.3390/jcm9061650 (2020).

17 Tseng, A. S., Crowell, M., Orenstein, R., Patron, R. L. & DiBaise, J. Older patient age is associated with similar safety but higher relapse after fecal microbiota transplantation for recurrent clostridium difficileInfection. Official journal of the American College of Gastroenterology | ACG 112, S51 (2017).

18 Reese, A. T. et al. Antibiotic-induced changes in the microbiota disrupt redox dynamics in the gut. Elife 7, doi:10.7554/eLife.35987 (2018).

19 Reese, A. T. et al. Microbial nitrogen limitation in the mammalian large intestine. Nat Microbiol 3, 1441–1450, doi:10.1038/s41564-018-0267-7 (2018).

20 Palleja, A. et al. Recovery of gut microbiota of healthy adults following antibiotic exposure. Nat Microbiol 3, 1255–1265, doi:10.1038/s41564-018-0257-9 (2018).

21 Ng, K. M. et al. Recovery of the gut microbiota after antibiotics depends on host diet, community context, and environmental reservoirs. Cell Host Microbe 28, 628, doi:10.1016/j.chom.2020.09.001 (2020).

22 Dethlefsen, L. & Relman, D. A. Incomplete recovery and individualized responses of the human distal gut microbiota to repeated antibiotic perturbation. Proc Natl Acad Sci U S A 108 Suppl 1, 4554–4561, doi:10.1073/pnas.1000087107 (2011).

23 Lavelle, A. et al. Baseline microbiota composition modulates antibiotic-mediated effects on the gut microbiota and host. Microbiome 7, 111, doi:10.1186/s40168-019-0725-3 (2019).

24 Chng, K. R. et al. Metagenome-wide association analysis identifies microbial determinants of post-antibiotic ecological recovery in the gut. Nat Ecol Evol 4, 1256–1267, doi:10.1038/s41559-020-1236-0 (2020).

25 Gibbons, S. M. Keystone taxa indispensable for microbiome recovery. Nat Microbiol 5, 1067–1068, doi:10.1038/s41564-020-0783-0 (2020).

26 Fujimoto, K. et al. Functional restoration of bacteriomes and viromes by fecal microbiota transplantation. Gastroenterology, doi:10.1053/j.gastro.2021.02.013 (2021).

27 Morgun, A. et al. Uncovering effects of antibiotics on the host and microbiota using transkingdom gene networks. Gut 64, 1732–1743, doi: 10.1136/gutjnl-2014-308820 (2015).

28 Ohkusa, T. [Production of experimental ulcerative colitis in hamsters by dextran sulfate sodium and changes in intestinal microflora]. Nihon Shokakibyo Gakkai Zasshi 82, 1327–1336 (1985).

29 Wirtz, S. et al. Chemically induced mouse models of acute and chronic intestinal inflammation. Nat Protoc 12, 1295–1309, doi:10.1038/nprot.2017.044 (2017).

30 Munyaka, P. M., Rabbi, M. F., Khafipour, E. & Ghia, J. E. Acute dextran sulfate sodium (DSS)-induced colitis promotes gut microbial dysbiosis in mice. J Basic Microbiol 56, 986–998, doi:10.1002/jobm.201500726 (2016).

31 Osaka, T. et al. Meta-Analysis of fecal microbiota and metabolites in experimental colitic mice during the inflammatory and healing phases. Nutrients 9, doi:10.3390/nu9121329 (2017).

32 De Fazio, L. et al. Longitudinal analysis of inflammation and microbiota dynamics in a model of mild chronic dextran sulfate sodium-induced colitis in mice. World J Gastroenterol 20, 2051–2061, doi:10.3748/wjg.v20.i8.2051 (2014).

33 Melgar, S., Karlsson, A. & Michaëlsson, E. Acute colitis induced by dextran sulfate sodium progresses to chronicity in C57BL/6 but not in BALB/c mice: correlation between symptoms and inflammation. Am J Physiol Gastrointest Liver Physiol 288, G1328–1338, doi:10.1152/ajpgi.00467.2004 (2005).

34 Tang, C. et al. Suppression of IL-17F, but not of IL-17A, provides protection against colitis by inducing T(reg) cells through modification of the intestinal microbiota. Nat Immunol 19, 755–765, doi:10.1038/s41590-018-0134-y (2018).

35 Cao, G. et al. Bacillus amyloliquefaciens ameliorates dextran sulfate sodium-induced colitis by improving gut microbial dysbiosis in mice model. Front Microbiol 9, 3260, doi:10.3389/fmicb.2018.03260 (2018).

36 Tabbaa, O. M., Aboelsoud, M. M. & Mattar, M. C. Long-term safety and efficacy of fecal microbiota transplantation in the treatment of Clostridium difficile infection in patients with and without inflammatory bowel disease: a tertiary care center’s experience. Gastroenterology Res 11, 397–403, doi:10.14740/gr1091 (2018).

37 Ding, X. et al. Long-term safety and efficacy of fecal microbiota transplant in active ulcerative colitis. Drug Saf 42, 869–880, doi:10.1007/s40264-019-00809-2 (2019).

38 Xiao, Y., Angulo, M. T., Lao, S., Weiss, S. T. & Liu, Y. Y. An ecological framework to understand the efficacy of fecal microbiota transplantation. Nat Commun 11, 3329, doi:10.1038/s41467-020-17180-x (2020).

39 Baruch, E. N. et al. Fecal microbiota transplant promotes response in immunotherapy-refractory melanoma patients. Science 371, 602–609, doi:10.1126/science.abb5920 (2021).

40 Davar, D. et al. Fecal microbiota transplant overcomes resistance to anti-PD-1 therapy in melanoma patients. Science 371, 595–602, doi:10.1126/science.abf3363 (2021).

41 Smith, P. et al. Regulation of life span by the gut microbiota in the short-lived African turquoise killifish. Elife 6, doi:10.7554/eLife.27014 (2017).

42 Bárcena, C. et al. Healthspan and lifespan extension by fecal microbiota transplantation into progeroid mice. Nat Med 25, 1234–1242, doi:10.1038/s41591-019-0504-5 (2019).

43 Stebegg, M. et al. Heterochronic faecal transplantation boosts gut germinal centres in aged mice. Nat Commun 10, 2443, doi:10.1038/s41467-019-10430-7 (2019).

44 D’Amato, A. et al. Faecal microbiota transplant from aged donor mice affects spatial learning and memory via modulating hippocampal synaptic plasticity- and neurotransmission-related proteins in young recipients. Microbiome 8, 140, doi:10.1186/s40168-020-00914-w (2020).

45 Kundu, P et al. Neurogenesis and prolongevity signaling in young germ-free mice transplanted with the gut microbiota of old mice. Sci Transl Med 11, doi:10.1126/scitranslmed.aau4760 (2019).

46 Kelly, C. P. & LaMont, J. T. Clostridium difficile--more difficult than ever. N Engl J Med 359, 1932–1940, doi:10.1056/NEJMra0707500 (2008).

47 Mamo, Y., Woodworth, M. H., Wang, T., Dhere, T. & Kraft, C. S. Durability and long-term clinical outcomes of fecal microbiota transplant treatment in patients with recurrent Clostridium difficile infection. Clin Infect Dis 66, 1705–1711, doi:10.1093/cid/cix1097 (2018).

48 He, Z. et al. Multiple fresh fecal microbiota transplants induces and maintains clinical remission in Crohn’s disease complicated with inflammatory mass. Sci Rep 7, 4753, doi:10.1038/s41598-017-04984-z (2017).

49 Li, P. et al. Timing for the second fecal microbiota transplantation to maintain the long-term benefit from the first treatment for Crohn’s disease. Appl Microbiol Biotechnol 103, 349–360, doi:10.1007/s00253-018-9447-x (2019).

50 Staley, C. et al. Durable long-term bacterial engraftment following encapsulated fecal microbiota transplantation to treat Clostridium difficile infection. mBio 10, doi:10.1128/mBio.01586-19 (2019).

51 Lahti, L., Salojärvi, J., Salonen, A., Scheffer, M. & de Vos, W. M. Tipping elements in the human intestinal ecosystem. Nat Commun 5, 4344, doi:10.1038/ncomms5344 (2014).

52 Van de Guchte, M. et al. Alternative stable states in the intestinal ecosystem: proof of concept in a rat model and a perspective of therapeutic implications. Microbiome 8, 153, doi:10.1186/s40168-020-00933-7 (2020).

53 Huang, S. et al. Human skin, oral, and gut microbiomes predict chronological age. mSystems 5, doi:10.1128/mSystems.00630-19 (2020).

54 Low, A., Soh, M., Miyake, S. & Seedorf, H. Host-age prediction from fecal microbiome composition in laboratory mice. bioRxiv (2020).

55 Littmann, E. R. et al. Host immunity modulates the efficacy of microbiota transplantation for treatment of Clostridioides difficile infection. Nat Commun 12, 755, doi:10.1038/s41467-020-20793-x (2021).

56 Chu, H. & Mazmanian, S. K. Innate immune recognition of the microbiota promotes host-microbial symbiosis. Nat Immunol 14, 668–675, doi:10.1038/ni.2635 (2013).

57 Zheng, D., Liwinski, T. & Elinav, E. Interaction between microbiota and immunity in health and disease. Cell Res 30, 492–506, doi:10.1038/s41422-020-0332-7 (2020).

58 Lai, Z. L. et al. Fecal microbiota transplantation confers beneficial metabolic effects of diet and exercise on diet-induced obese mice. Sci Rep 8, 15625, doi:10.1038/s41598-018-33893-y (2018).

59 Tan, Y., Tian, Y., Chen, J., Yin, Z. & Yang, H. DPSN: standardizing the short names of amplicon-sequencing primers to avoid ambiguity. bioRxiv (2020).

60 Bolyen, E. et al. Reproducible, interactive, scalable and extensible microbiome data science using QIIME 2. Nat Biotechnol 37, 852–857, doi:10.1038/s41587-019-0209-9 (2019).

61 Martin, M. Cutadapt removes adapter sequences from high-throughput sequencing reads. EMBnet. journal 17, 10–12 (2011).

62 Callahan, B. J. et al. DADA2: High-resolution sample inference from Illumina amplicon data. Nat Methods 13, 581–583, doi:10.1038/nmeth.3869 (2016).

63 Price, M. N., Dehal, P. S. & Arkin, A. P. FastTree 2--approximately maximum-likelihood trees for large alignments. PLoS One 5, e9490, doi:10.1371/journal.pone.0009490 (2010).

64 Bokulich, N. A. et al. Optimizing taxonomic classification of marker-gene amplicon sequences with QIIME 2’s q2-feature-classifier plugin. Microbiome 6, 90, doi:10.1186/s40168-018-0470-z (2018).

65 Segata, N. et al. Metagenomic biomarker discovery and explanation. Genome Biol 12, R60, doi:10.1186/gb-2011-12-6-r60 (2011).

66 Bolger, A. M., Lohse, M. & Usadel, B. Trimmomatic: a flexible trimmer for Illumina sequence data. Bioinformatics 30, 2114–2120, doi:10.1093/bioinformatics/btu170 (2014).

67 Wood, D. E., Lu, J. & Langmead, B. Improved metagenomic analysis with Kraken 2. Genome Biol 20, 257, doi:10.1186/s13059-019-1891-0 (2019).

68 Franzosa, E. A. et al. Species-level functional profiling of metagenomes and metatranscriptomes. Nat Methods 15, 962–968, doi:10.1038/s41592-018-0176-y (2018).

69 Dobin, A. et al. STAR: ultrafast universal RNA-seq aligner. Bioinformatics 29, 15–21, doi:10.1093/bioinformatics/bts635 (2013).

70 Love, M. I., Huber, W. & Anders, S. Moderated estimation of fold change and dispersion for RNA-seq data with DESeq2. Genome Biol 15, 550, doi:10.1186/s13059-014-0550-8 (2014).

71 Wickham, H. (2016). ggplot2: Elegant graphics for data analysis. (Springer-Verlag, New York, USA), p. 260. ISBN 978-3-319-24277-4.

