## Supplement figures S1-S8 for "Establishment and Resilience of Transplanted Gut Microbiota in Aged Mice"

**Supplementary figures**

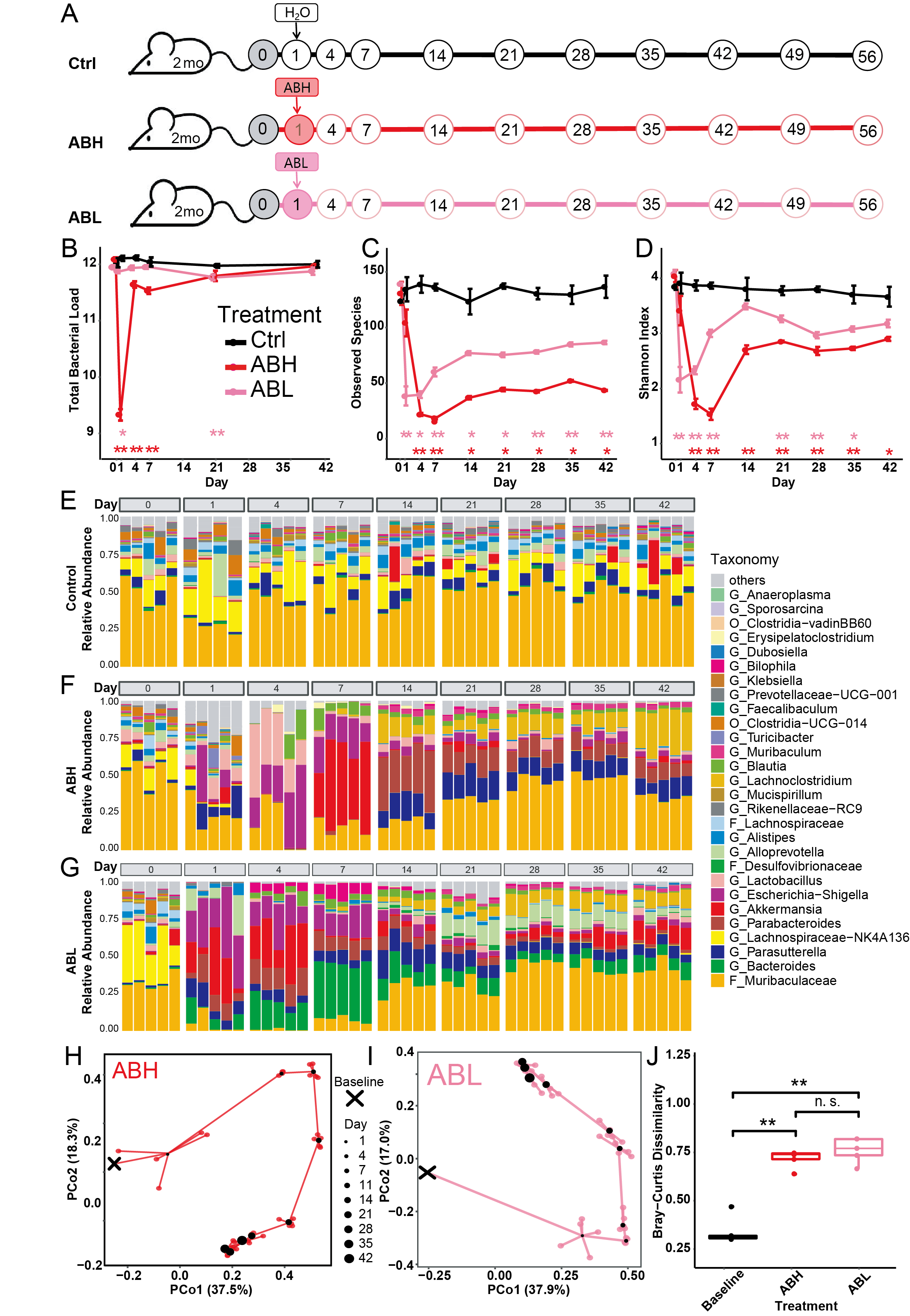

**Figure S1. Spontaneous recovery of gut microbiota following antibiotics treatment in young mice, related to Figure 1.** (**A**) Young mice were treated with the antibiotic cocktail on Day 1. Ctrl, Control; ABH, antibiotics-high dose; ABL, antibiotics-low dose; 20mo: 20-month-old mice. Circles are marked by time points (Day) of fecal sample collection. (**B**) Total microbial load (16S copy number log/gram) in fecal samples assayed by qPCR. (**C-D**) The number of observed species (**C**) and the Shannon diversity index (**D**) did not recover to the baseline level. Colored asterisks indicate statistically significant difference between the experimental groups and the control group. Error bar= SEM. (**E-G**) Changes in the relative abundance of microbial taxa during spontaneous recovery. O, order, F, family, G, genus. (**H-I**) Principal Coordinate Analysis (PCoA) based on Bray-Curtis dissimilarity, showing the trajectory of microbiota composition for ABH (**H**) and ABL (**I**) groups. Baseline (black cross) indicates the microbiota composition averaged over young mice in the control group from Day 1 to Day 56. For each time point, the central black dot indicates the average of 5 mice and the colored lines connect replicate samples. (**J**) Bray-Curtis dissimilarity between the gut microbiota composition on Day 56 and the Baseline composition. N=5 for each time points. The FDR-adjusted P values are shown between various treatment groups (two-sided Wilcoxon signed-rank test). **P* < 0.05 ***P*<0.01, ****P*<0.001. Some differences were observed between the restoration process in young and aged mice. For example, there were no difference in the effects on microbiota diversity between high-dose and low-dose antibiotics in aged mice (**Figure 1C-D**). By contrast, in young mice, the reduction and recovery of microbiota diversity in ABH and ABL groups were different (**Figure S1C-D**). However, in young mice the gut microbiota after spontaneous recovery showed no differences between ABH and ABL groups on Day 56 (**Figure S1G**), whereas in aged mice the difference is significant (**Figure 1G**).
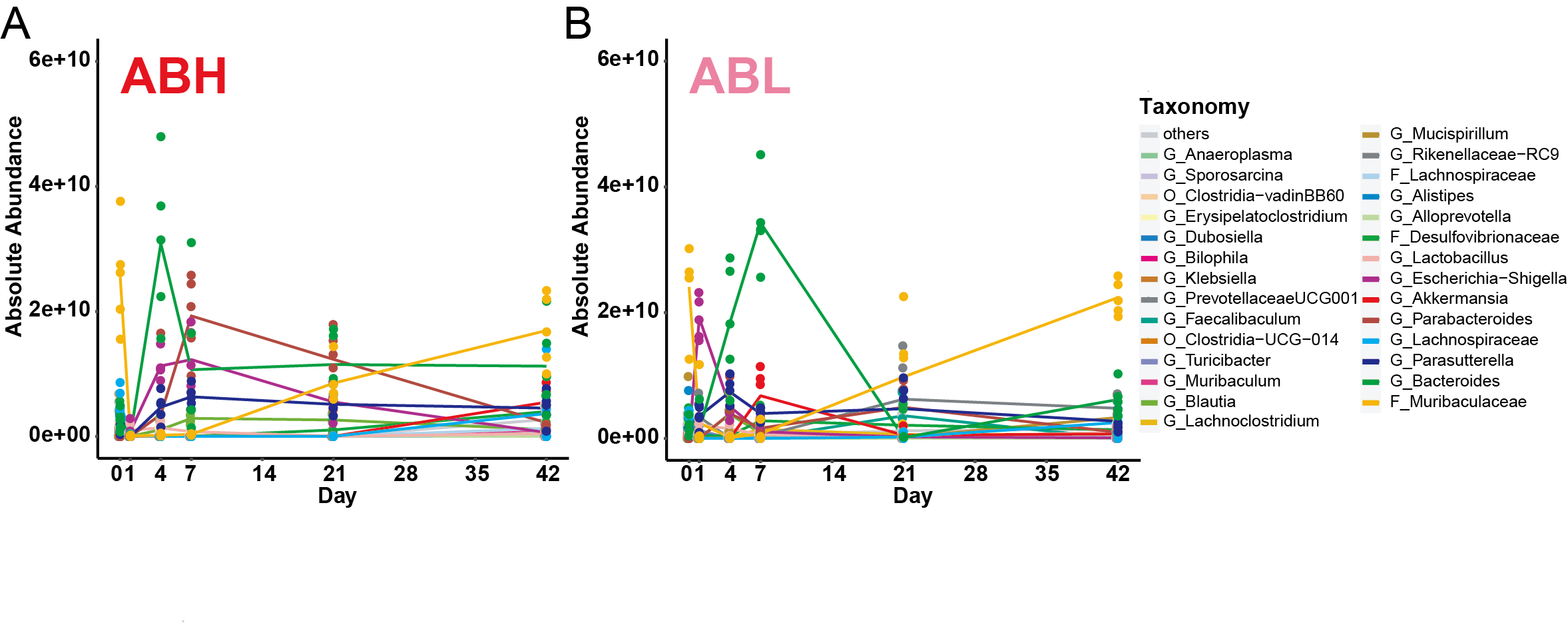

**Figure S2. Changes in the absolute abundance of bacteria taxa following antibiotics treatment, related to Figure 1.** The absolute abundance of bacteria taxa in the ABH (high-dose antibiotics) (**A**) and ABL (low-dose antibiotics) (**B**) groups. The absolute abundance was calculated by multiplying the relative abundance of each bacterial taxa with the total bacteria load (see Methods). A recent investigation by Chng et al. showed that the microbiota recovery after antibiotics treatment followed a “secondary succession” pattern in which primary species degrading complex substrates like polysaccharide grows fast and enables the increases of secondary species that produces short-chain fatty acid, followed by the final increase of tertiary species that promotes the diversity of the microbiota during the later stages of succession^25,26^. Based on the conditional co-occurrence analysis, the investigators identified some taxa as “recovery associated bacterial (RAB) species” whose presence were indispensable for the recovery. Some of the RAB species were also seen in our present study of aged and young mice, such as an early increase in *Bacteroides* and *Alistipes* (**Figure 1, Figure S1-S2**). The absolute abundance of *Bacteroides* briefly dominated the gut microbiota of mice in the ABH (or ABL) group on Day 4 (or Day 7, respectively), then quickly decreased and stabilized. Other taxa, e.g., *Escherichia-Shigella, Parasutterella, Parabacteroides* and *Akkermansia*, also showed similar “transient peaks” in their absolute abundances (**Figure S2**). Most of these peaks appeared during Day 1- Day 7, suggesting that the first week after antibiotic treatment may be a critical time window for microbiota recovery. Further studies are needed to understand the interaction network between the host environment and the RAB species during the ecological succession following antibiotics treatment.

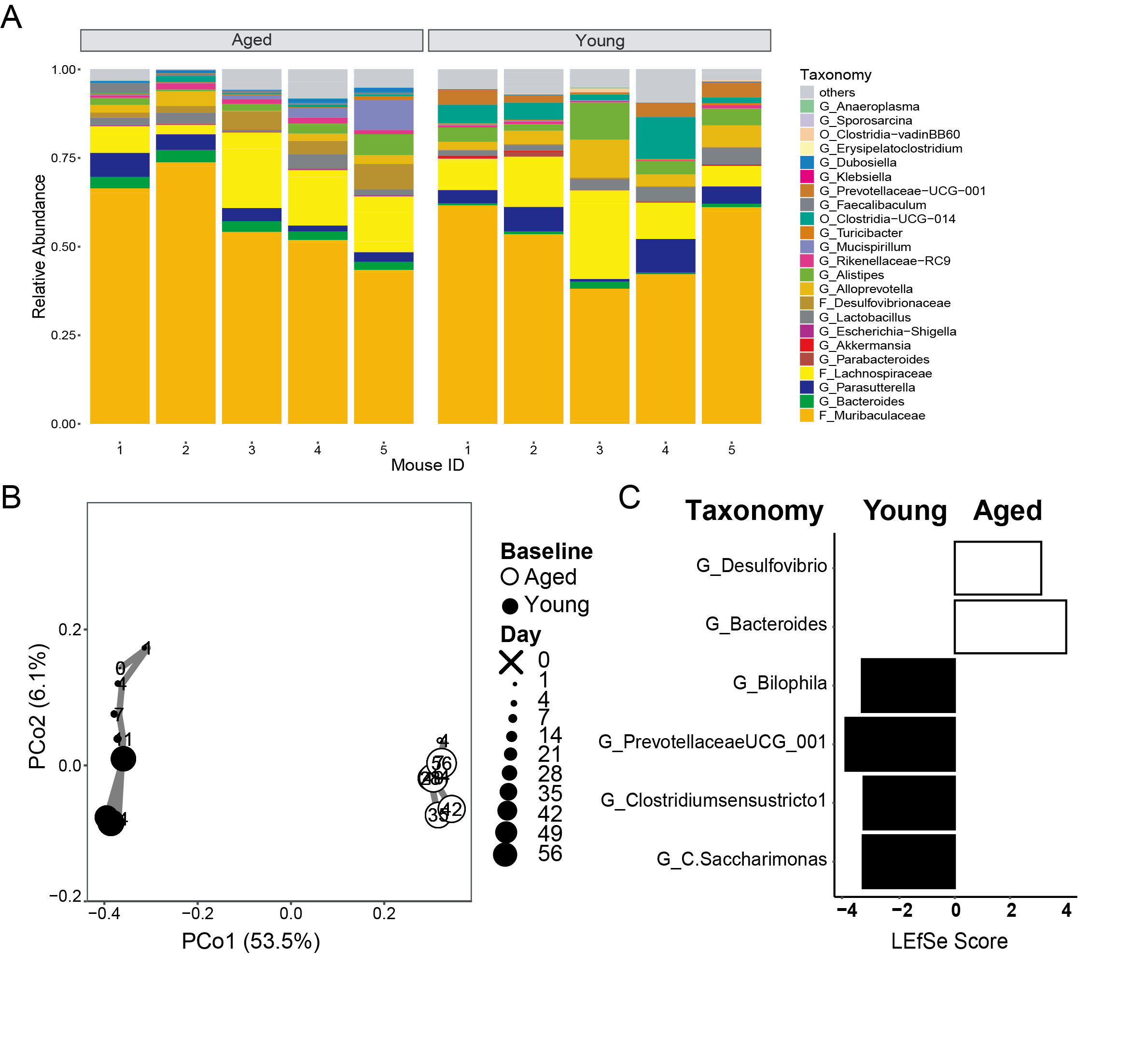

**Figure S3. The baseline gut microbiota of young and aged mice, related to Figure 2.** (**A**) stacking bar plots showing the relative abundance of microbial taxa in young and aged mice. O, order, F, family, G, genus. (**B**) Principal coordinate analysis based on Bray-Curtis dissimilarities of gut microbiota composition of young mice (2-month-old) and aged mice (20-month-old). Aged mice were used as matched donors for FMT (FMT-M group). Young mice were used as unmatched donors for FMT (FMT-UM group). N=5 for each group at each time point. There was a minor shift in the gut microbiota composition of young mice, most likely due to the natural aging process. Overall, the gut microbiota composition of young and aged mice was found to be stable over 6-8 weeks (**Figure 1E, Figure S1E**). (**C**) Bacterial taxa enriched in the gut microbiota of young mice (black) versus aged mice (white) were identified by LEfSe analysis.

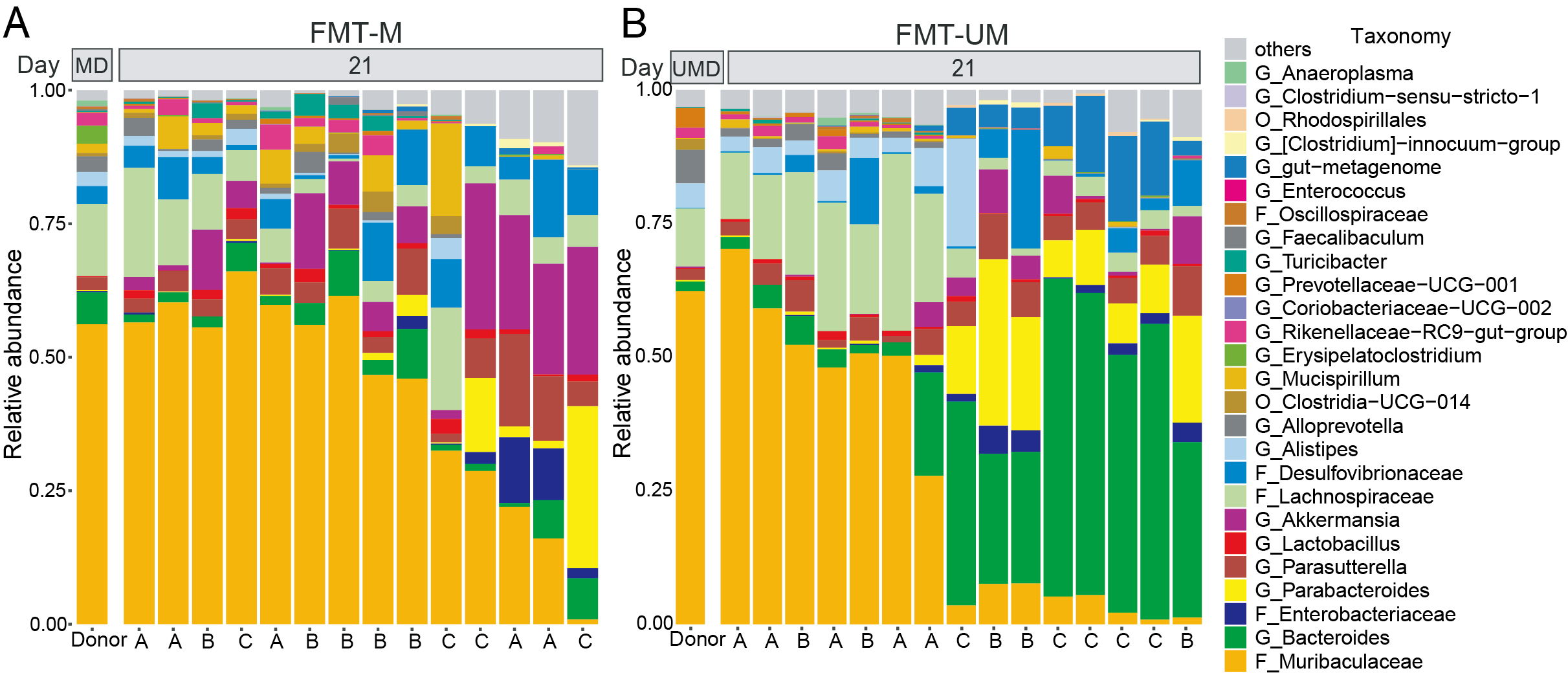

**Figure S4.** **The gut microbiota compositions of FMT-M and FMT-UM groups on Day 21, related to Figure 2.** The relative abundance of gut microbial taxa in aged mice received FMT-M (**A**) or FMT-UM (**B**) on day 21. The first column is the average composition of donor samples: MD, matched donors; UMD, unmatched donors. Samples are ordered based on their Bray-Curtis dissimilarity to the donor. Mice housed in the same cage (3 ages labelled as A, B, C) are indicated at the bottom of each column. The difference among individual gut microbiota is not fully attributable to the cage effect.

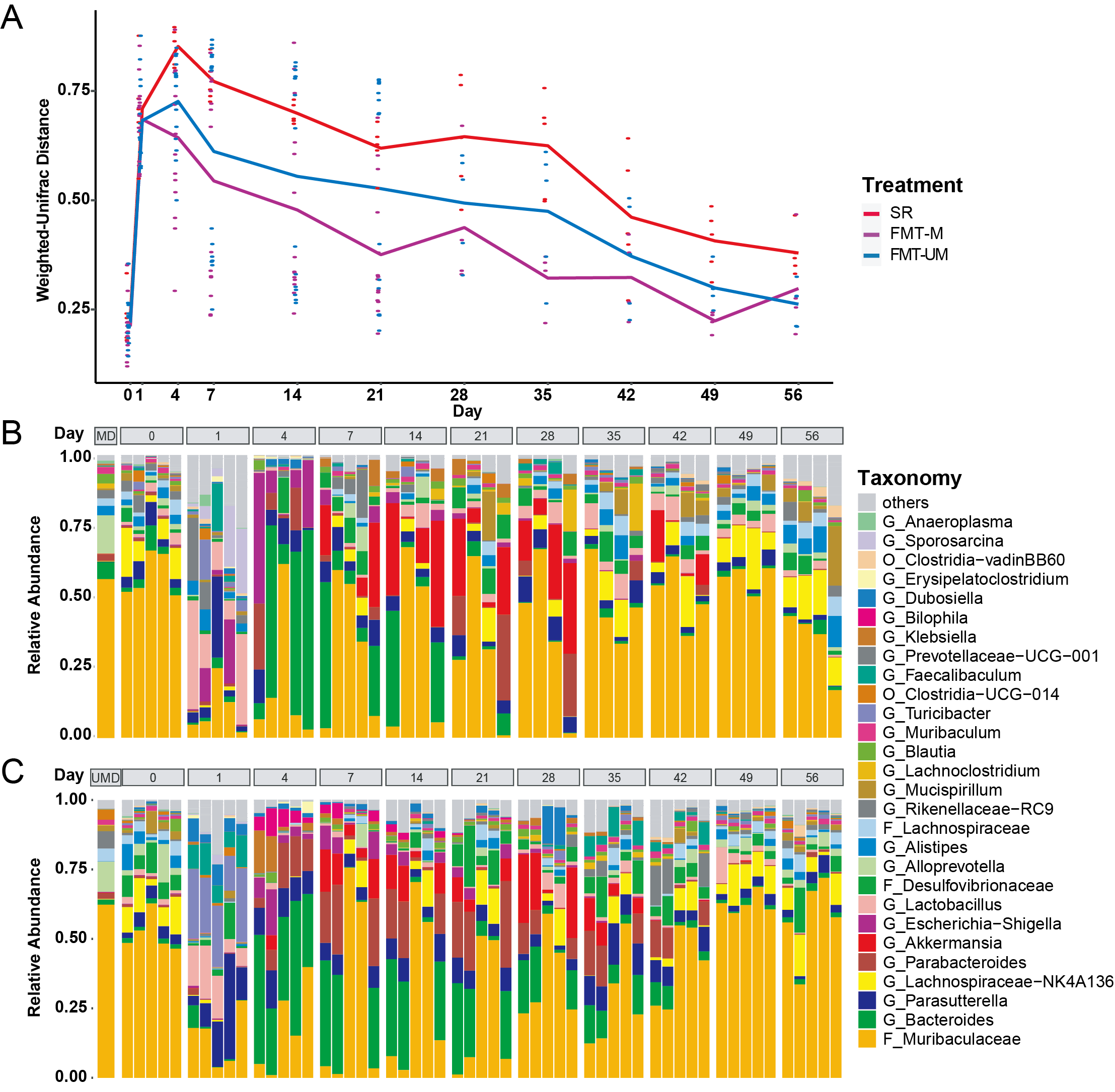

**Figure S5.** **The gut microbiota compositions throughout the long-term recovery, related to Figure 2.** (**A**) The weighted Unifrac distance between the gut microbiota of treatment groups and the baseline microbiota of aged mice throughout the long-term recovery. For FMT-M and FMT-UM groups, N=15 until Day 21 and N=5 afterwards. N=5 for SR group. (**B-C**) The relative abundance of gut microbial taxa in aged mice received FMT-M (**B**) or FMT-UM (**C**)**.** The first column is the average composition of donor samples: MD, matched donors; UMD, unmatched donors.

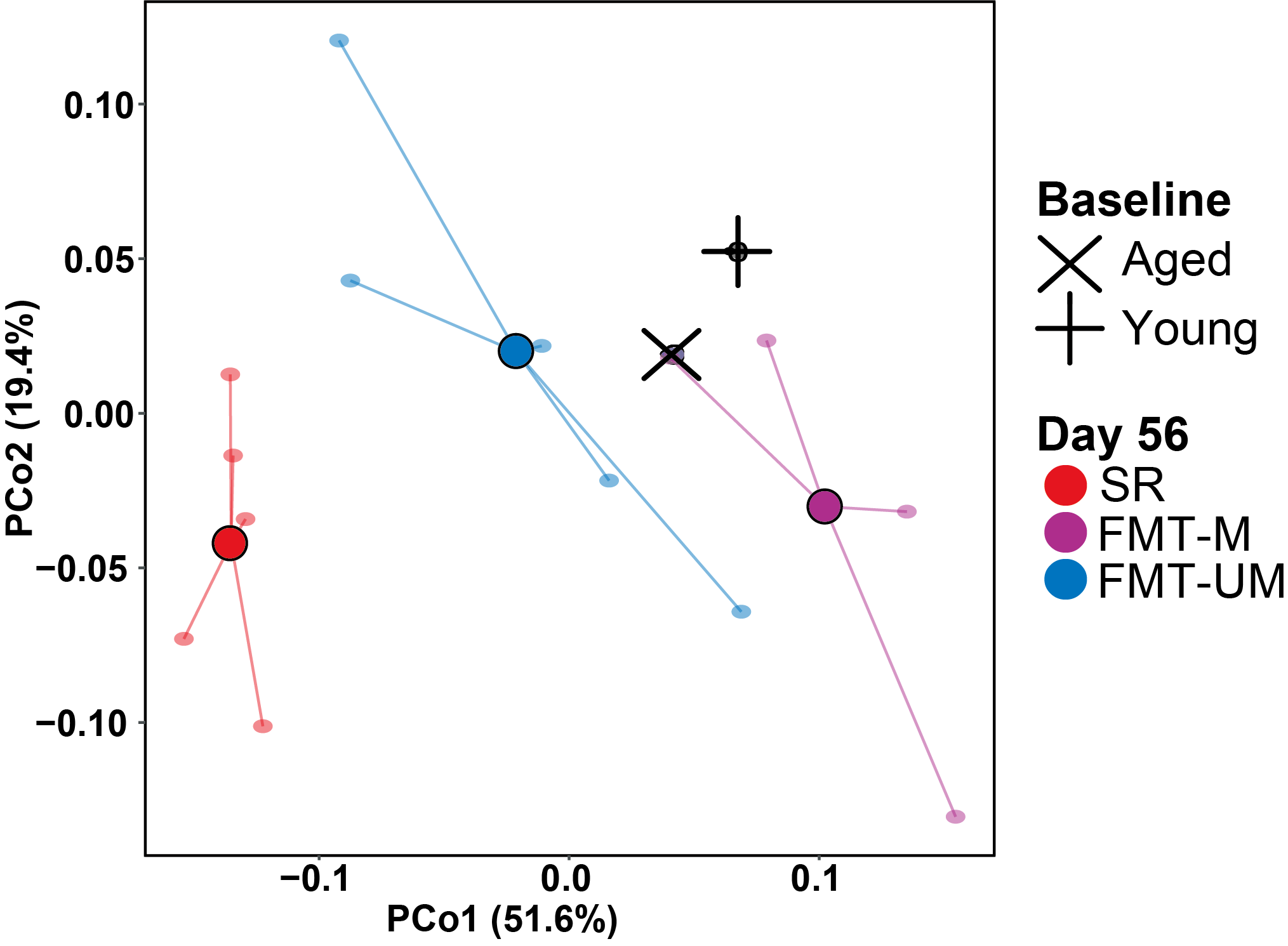

**Figure S6. Principal coordinate analysis of the abundance of metagenomic gene pathways.** For each group, the central dot indicates the mean (N=5) and the lines connect individual samples.

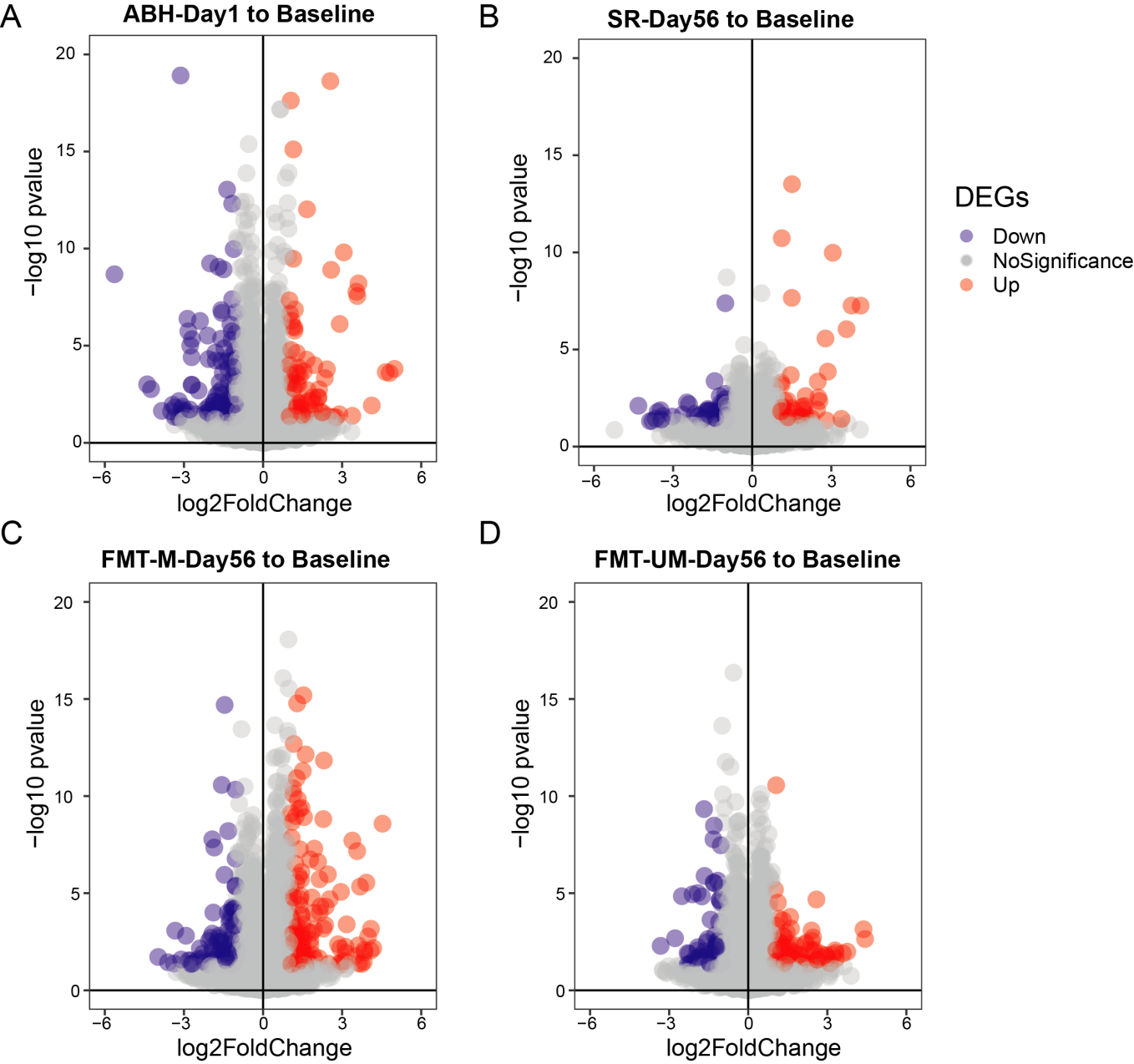

**Figure S7. Volcano plots of DEGs (*P* < 0.05) between each group and aged baseline found in RNA sequencing.**

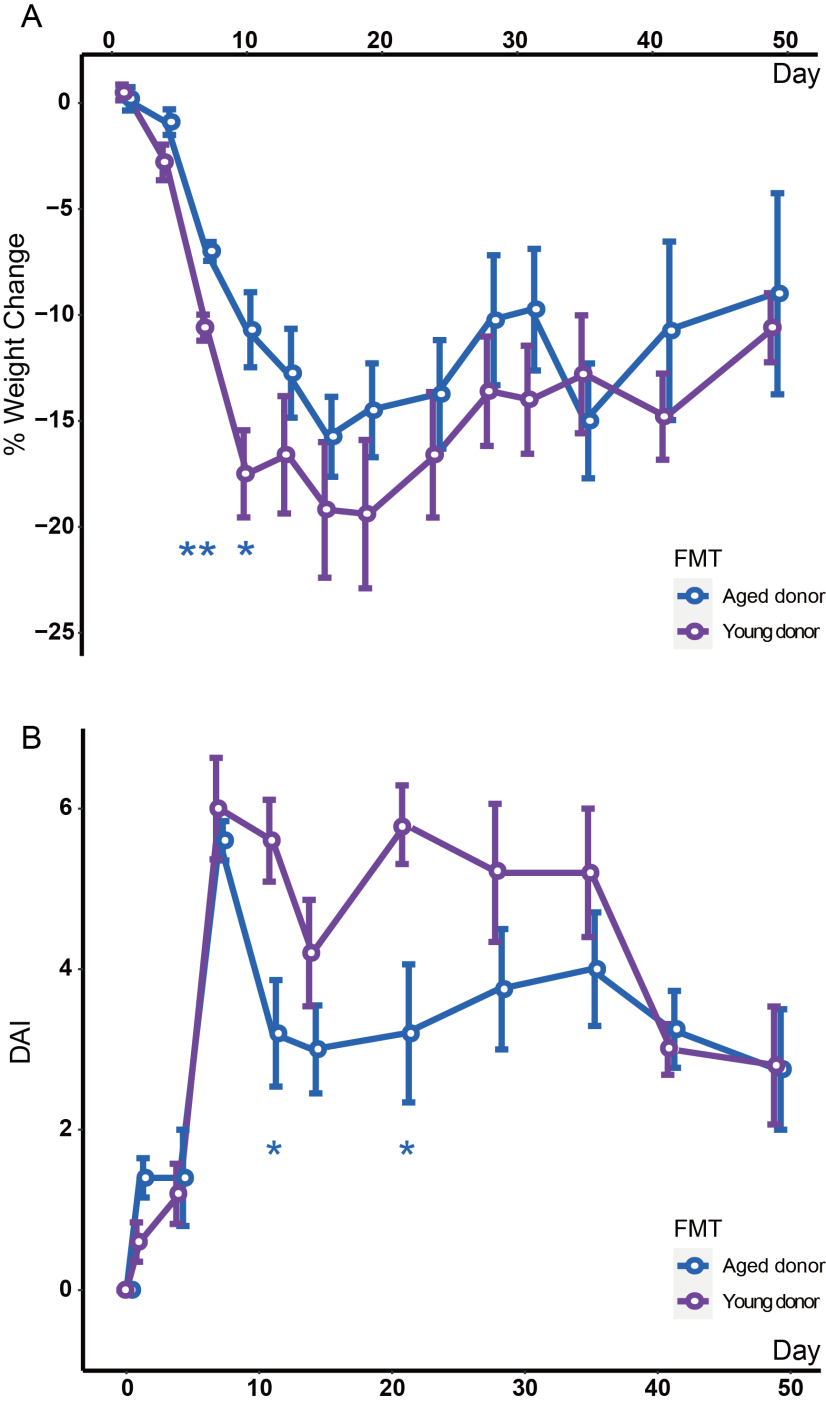

**Figure S8. DSS treatment induce acute and self-limiting colitis symptoms in aged recipients of FMT-M or FMT-UM.** (**A**) weight changes. (**B**) disease activity index. N=4 (due to the death of one mouse) for FMT-M and N=5 for FMT-UM group on each time point. Error bar= SEM. *indicates significant difference at *P* < 0.05, **, *P*<0.01, ***, *P*<0.001 analyzed by FDR-adjusted *P* values (two-sided Wilcoxon signed-rank test).
